# *Bacillus subtilis* RNase HII is inefficient at processing guanosine monophosphate and damaged ribonucleotides

**DOI:** 10.1101/2025.09.17.676834

**Authors:** Julianna R. Cresti, Lyle A. Simmons

## Abstract

During one round of DNA replication, nearly 2,000 ribonucleoside monophosphates (rNMPs) are incorporated in place of their cognate deoxyribonucleoside monophosphate (dNMP). Given their high rate of insertion, genomic DNA would contain rNMPs that are damaged or mismatched. Here, we tested the activity of *Bacillus subtilis* and *Escherichia coli* RNase HII on all four canonical, mismatched, and damaged rNMPs. We show that *E. coli* RNase HII is adept at incising most rNMP variants from DNA at similar frequencies, with the exception of an oxidized rNMP, where endoribonuclease activity is sharply reduced. In contrast, *B. subtilis* RNase HII efficiently incised rAMP, rCMP, and rUMP, but was inefficient at processing rGMP in both a canonical and mismatched base pair. We tested damaged ribonucleotides and found that *B. subtilis* RNase HII is refractory to processing abasic and oxidized ribonucleotide lesions. Our work shows that bacterial RNase HII enzymes have different intrinsic endoribonuclease activity toward the repair of canonical, mismatched, and damaged rNMPs, demonstrating that not all rNMP errors provoke efficient resolution. Our finding that *B. subtilis* RNase HII is recalcitrant to repairing damaged rNMPs resembles what is observed for eukaryotic RNase H2 orthologs, suggesting that other repair processes are necessary to resolve damaged rNMPs.

## INTRODUCTION

Ribonucleotides are routinely incorporated into genomic DNA through the essential process of DNA replication (Nick McElhinny *et al*., 2010a, Yao *et al*., 2013). It is well established that bacterial and eukaryotic replicative DNA polymerases make errors by incorporating a ribonucleoside monophosphate (rNMP) in the place of its corresponding deoxyribonucleoside monophosphate (dNMP) (Yao *et al*., 2013, Schroeder *et al*., 2017, Nick McElhinny *et al*., 2010a, Nick McElhinny *et al*., 2010b). In bacteria, it has been reported that rNMP errors occur *in vitro* about 2,000 times per round of replication (Yao *et al*., 2013). These errors have been measured *in vivo* with numbers ranging between about 200 to over 2,200 per round of replication across different bacterial species (Zatopek *et al*., 2019, Kouzminova *et al*., 2017). Similarly, in eukaryotes, rNMP errors in *Saccharomyces cerevisiae* can exceed 10,000 per round of replication and have been predicted to be in the millions for multicellular eukaryotes, including mice and humans (Nick McElhinny *et al*., 2010b, Williams & Kunkel, 2014, Zhou *et al*., 2021). These errors are often referred to as “sugar errors” because, although the nitrogenous base is correct, the ribose sugar has a hydroxyl group (-OH) at the C2′ position instead of a hydrogen atom. The 2′-OH is reactive, with the potential to become a good nucleophile, resulting in phosphodiester bond cleavage (Oivanen *et al*., 1998), forming a cyclic 2′, 3′-phosphate that cannot be ligated or act as a substrate for extension by a DNA polymerase (Oivanen *et al*., 1998). Further, ribonucleotides can be mutagenic upon removal. rNMPs can be removed by topoisomerase I in eukaryotes, but this causes single-stranded breaks and deletions (Kim *et al*., 2011). Likewise, in bacteria, rNMP errors can be removed using an error-prone pathway that results in GC-AT transitions (Schroeder *et al*., 2017). Given the propensity for single ribonucleotide errors to occur *in vivo*, it has been estimated that rNMPs represent the most frequent type of nucleotide in need of repair in genomes ranging from bacteria to humans (Nick McElhinny *et al*., 2010b, Schroeder *et al*., 2015, Williams *et al*., 2016, Zhou *et al*., 2021).

Ribonucleotides embedded in the genome are repaired through ribonucleotide excision repair (RER), which has been previously studied in *Escherichia coli*, archaea, yeast, and multicellular eukaryotes (Sparks *et al*., 2012, Schroeder *et al*., 2017, Heider *et al*., 2017). Type 2 Ribonuclease H (RNase H) enzymes, Ribonuclease HII (RNase HII) in bacteria and archaea or Ribonuclease H2 (RNase H2) in eukaryotes, initiate this process. RNase HII binds the ribonucleotide in double-stranded DNA (dsDNA) and hydrolyzes the phosphodiester bond 5′ to the ribonucleotide (Itaya *et al*., 1999, Ohtani *et al*., 1999a). Acidic residues, together known as a DEDD motif, coordinate divalent metal cations, preferentially Mg^2+^, and facilitate hydrolysis of the backbone (Rychlik *et al*., 2010, Randall *et al*., 2017). During the canonical RER pathway in bacteria, DNA polymerase I binds the nick, removes the ribonucleotide using its 5′-3′ exonuclease activity, and substitutes the rNMP with the corresponding dNMP (Schroeder *et al*., 2017). RNase H2 in eukaryotes also incises the dsDNA 5′ to the rNMP (Rychlik *et al*., 2010). The rNMP is then removed by flap endonuclease 1 (FEN1), while DNA polymerase δ (Pol δ), aided by proliferating cell nuclear antigen (PCNA) and the clamp loader, catalyze strand displacement synthesis (Sparks *et al*., 2012). Since rNMPs are incorporated at such a high frequency, redundancy in this pathway occurs — Exo1 can function in place of FEN1, and DNA polymerase ε can replace Pol δ during strand displacement synthesis and gap filling (Sparks *et al*., 2012). In bacteria, the redundancies are less clear. There is evidence in *E. coli* that nucleotide excision repair (NER) can function in the absence of RNase HII and translesion DNA polymerases can participate in the resynthesis step, albeit increasing mutagenesis (Vaisman *et al*., 2013, Vaisman *et al*., 2014). Further, additional results in *E. coli* show that the surveillance and recognition of ribonucleotides by RNase HII is coupled to RNA polymerase (RNAP), showing that RNase HII is physically associated with RNAP, allowing for ribonucleotides to be efficiently detected in DNA undergoing transcription (Hao *et al*., 2023).

Due to the sheer number of ribonucleotides incorporated into genomic DNA, it is expected that some rNMPs will become damaged or otherwise modified (Randerath *et al*., 1992, Loeb & Preston, 1986, Cheng *et al*., 1992, Malfatti *et al*., 2017). However, it remains unclear if and how cells can repair mismatched or damaged ribonucleotides. Prior work has shown that replicative DNA polymerases are poor at using their 3′-5′ “proofreading” exonuclease activity to remove properly base-paired ribonucleotides (Williams *et al*., 2012). When rNMPs are paired with the incorrect complement and form a mismatch, they can serve as a target for both mismatch repair (MMR) by MutSα and RNase H2, as shown in *S. cerevisiae* (Shen *et al*., 2011). *E. coli* RNase HII has been shown to incise mismatched rNMPs, while *Pyrococcus abyssi* RNase HII was shown to be unable to incise mismatched rNMPs by one group and described as less efficient to incise mismatched rNMPs in another study (Malfatti *et al*., 2019, Reveil *et al*., 2023). Therefore, on balance, most type 2 RNase H enzymes are at least capable of endoribonuclease activity on mismatched rNMPs, while some enzymes, like *E. coli* RNase HII, have been shown to prefer mismatched rNMP substrates (Malfatti *et al*., 2019). Moreover, given the high rate of misincorporation, damaged rNMPs, including abasic or AP sites (apurinic and apyrimidinic, rOH) and oxidative damage, including 8-oxoguanosine (r8oG), would be expected to occur *in vivo* (Malfatti *et al*., 2017, Malfatti *et al*., 2019). Interestingly, rOH and r8oG do not serve as substrates for eukaryotic or archaeal RNase H2 (Malfatti *et al*., 2017, Malfatti *et al*., 2019). In contrast, *E. coli* RNase HII has been reported to incise both lesions *in vitro* (Malfatti *et al*., 2019). To our knowledge, since *E. coli* RNase HII is the only bacterial type 2 RNase H enzyme tested on mismatched and damaged rNMPs, it remains unclear if other bacterial RNase HIIs are capable of processing rNMP variants.

In this study, we compare the activity of *E. coli* and *Bacillus subtilis* RNase HII on all four properly base-paired rNMPs, mismatched rNMPs, and damaged rNMPs. We find that *E. coli* RNase HII is efficient at resolving properly base-paired rAMP, rGMP, rCMP, and rUMP, as well as mismatched rNMPs, but shows no or weak activity on damaged rNMP lesions. Strikingly, while *B. subtilis* RNase HII efficiently cleaves rAMP, rUMP, and rCMP, the enzyme demonstrates poor processing of rGMP in a Watson-Crick base pair or mismatched context. We find that *B. subtilis* RNase HII is refractory to incision at rOH and r8oG, suggesting a substrate preference that more closely resembles that of eukaryotic RNase H2 than *E. coli* RNase HII. Because *B. subtilis* RNase HII is inefficient at processing guanosine monophosphate, we suggest that differences in bacterial RNase HII substrate specificity will shape genome stability through differences in processing canonical, mismatched and damaged rNMPs *in vivo*. Further, given that the bacterial RNase HII enzymes tested herein are either slow or unable to process damaged rNMPs, we conclude that damaged rNMPs are likely to be repaired through the appropriate base excision repair (BER) pathway as opposed to RER in many bacteria.

## RESULTS

### EcoRNase HII efficiently incises all four canonical rNMPs

*E. coli* RNase HII (EcoRNase HII) is the most well-studied bacterial RNase HII enzyme (Friedberg *et al*., 2006). EcoRNase HII has been tested for activity on a variety of rNMP-containing substrates *in vitro* (Ohtani *et al*., 2000, Rydberg & Game, 2002, Malfatti *et al*., 2019, Friedberg *et al*., 2006). To our knowledge, EcoRNase HII activity has not yet been compared across all four canonical rNMPs in the same study (Itaya, 1990, Ohtani *et al*., 2000, Haruki *et al*., 2002). We used 25-oligonucleotide dsDNA containing rA:dT, rG:dC, rC:dG, and rU:dA. Each substrate was incubated with EcoRNase HII at 37°C, followed by electrophoresis using denaturing urea-PAGE to resolve the fluorophore-labeled rNMP-containing strand. For assessment of each rNMP, we included a no protein control, a single-stranded DNA (ssDNA) containing the rNMP, and a catalytically impaired RNase HII protein variant (D16A, E17A). Substrate was treated with NaOH to induce alkaline hydrolysis and generate a marker for incision. In the case of EcoRNase HII, rAMP, rGMP, rCMP, and rUMP show similar percentages of incision, with the reaction about 50% complete by 1 minute and near 100% complete by 5 minutes **(Figure 1)**. EcoRNase HII does appear slightly less efficient on rCMP with ∼32% incised by 1 minute **(Figure 1C)**. Although this difference was not statistically significant when compared to rAMP. We conclude that EcoRNase HII incises all four rNMPs with comparable efficiencies. Of note, we show that EcoRNase HII efficiently incises rUMP. Uracil DNA glycosylases (UDGs), which are involved in BER, recognize dUMP in DNA (Hayakawa *et al*., 1978, Lindahl, 1974). UDGs can remove uracil from both ssDNA and dsDNA, but not ssRNA due to steric clash of a conserved phenylalanine residue with the 2′-OH (Pearl, 2000, Savva *et al*., 1995). Given that UDG is unlikely to act upon a ribonucleotide substrate, we conclude that RER is responsible for repairing rUMP in genomic DNA (see discussion).

**Figure 1.**
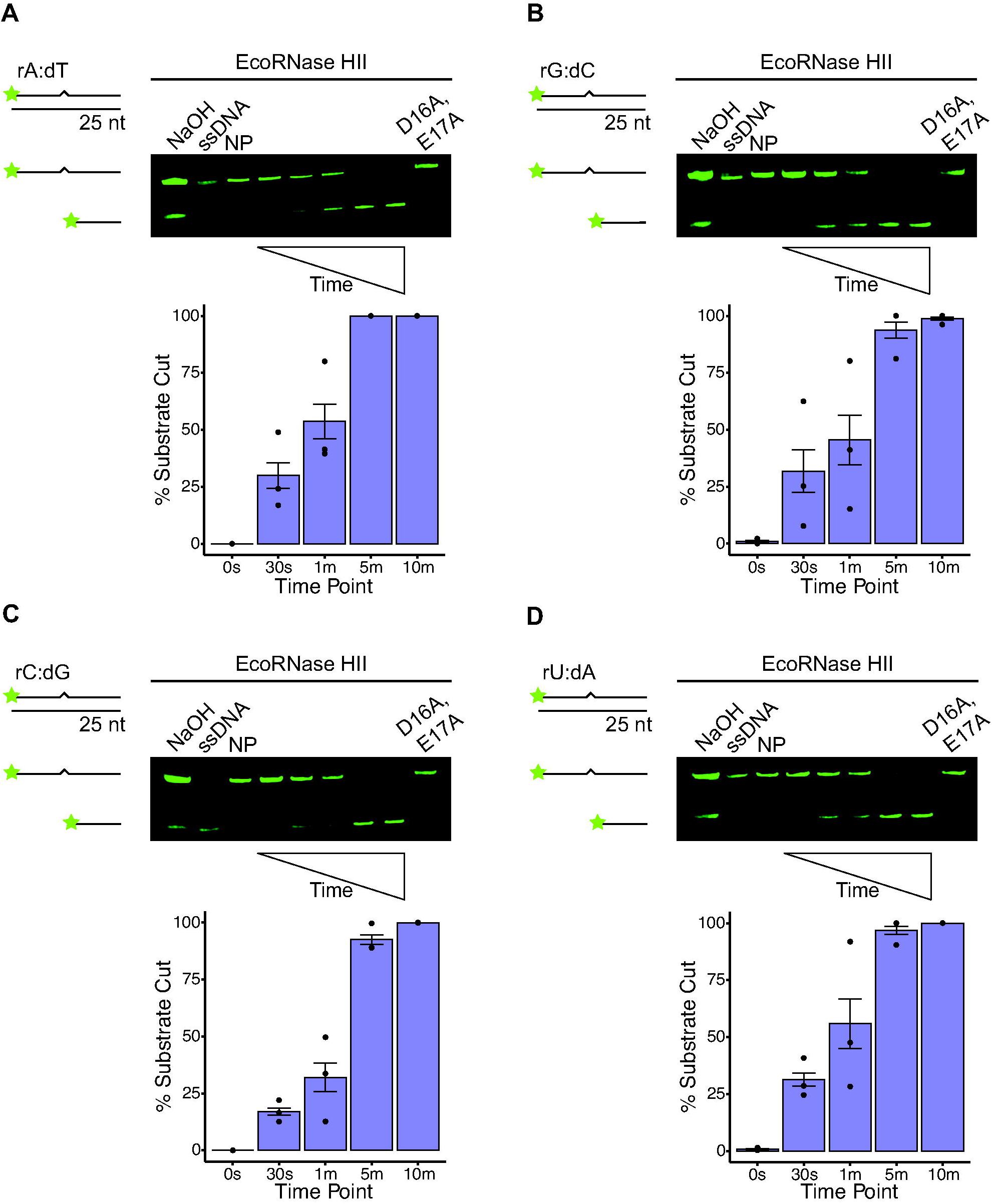
EcoRNase HII demonstrates similar nuclease activity on all four single rNMPs embedded in DNA. Representative urea-PAGE for assays performed over 10 minutes at 37°C with 6.25 nM *E. coli* RNase HII and 100 nM of the following 25-nucleotide substrates: (A) rA:dT, (B) rG:dC, (C) rC:dG, and (D) rU:dA. Each substrate is composed of 25-nucleotide long dsDNA with a single rNMP, represented by a zigzag. Quantification below each gel shows the mean percent of substrate incised over time (0 s, 30 s, 1 min., 5 min., and 10 min. time points) with black bars to show the standard error, for three replicates. Each gel contains ssDNA, no protein (NP), and catalytic impaired (D16A, E17A) controls. An alkaline ladder, represented by “NaOH”, was prepared by incubating substrate with 200 nM final NaOH. rA:dT was prepared with oJC3 and oJC2, rG:dC was prepared with oJC3 and oJC5, rC:dG was prepared with oJC6 and oJC7, and rU:dA was prepared with oJC8 and oJC9. Any difference observed for incision was not statistically significant.

### BsuRNase HII inefficiently incises rG:dC

We followed the same assay procedure as described for EcoRNase HII but instead with *B. subtilis* RNase HII (BsuRNase HII). Like EcoRNase HII, BsuRNase HII has not been tested on all four canonical rNMPs in the same study (Itaya *et al*., 1999, Randall *et al*., 2017). Both EcoRNase HII and BsuRNase HII were tested in the same manner, although we used 50 nM BsuRNase HII as compared with 6.25 nM EcoRNase HII to achieve a similar percent of enzyme activity. Although there are moderate differences, we generally found that BsuRNase HII incised rAMP, rCMP and rUMP with similar efficiencies, as each reaction reached ∼14-24% incision by 1 minute **(Figure 2A, C, and D)**. We compared the percent incision of each time point to rAMP for all other substrates. We found a statistically significant increase in rCMP and rUMP incision for several time points. We found a statistically significant decrease in rGMP incision at three of the four time points collected. Thus, BsuRNase HII was very slow to incise rGMP, showing only ∼0.5% incision by 1 minute and <50% incision by 5 minutes **(Figure 2B)**. Slow incision at rGMP was consistent among all the replicates and conditions tested over the course of this study. As controls, we used no protein, a ssDNA containing the rNMP, and a catalytically-impaired BsuRNase HII variant (D78A, E79A). An alkaline ladder was generated by incubating substrate with NaOH to induce hydrolysis. We conclude that BsuRNase HII is inefficient at cleaving rGMP in DNA, and like EcoRNase HII, is capable of efficient removal of rUMP from DNA, further supporting the conclusion that rUMP is primarily repaired by RER.

**Figure 2.**
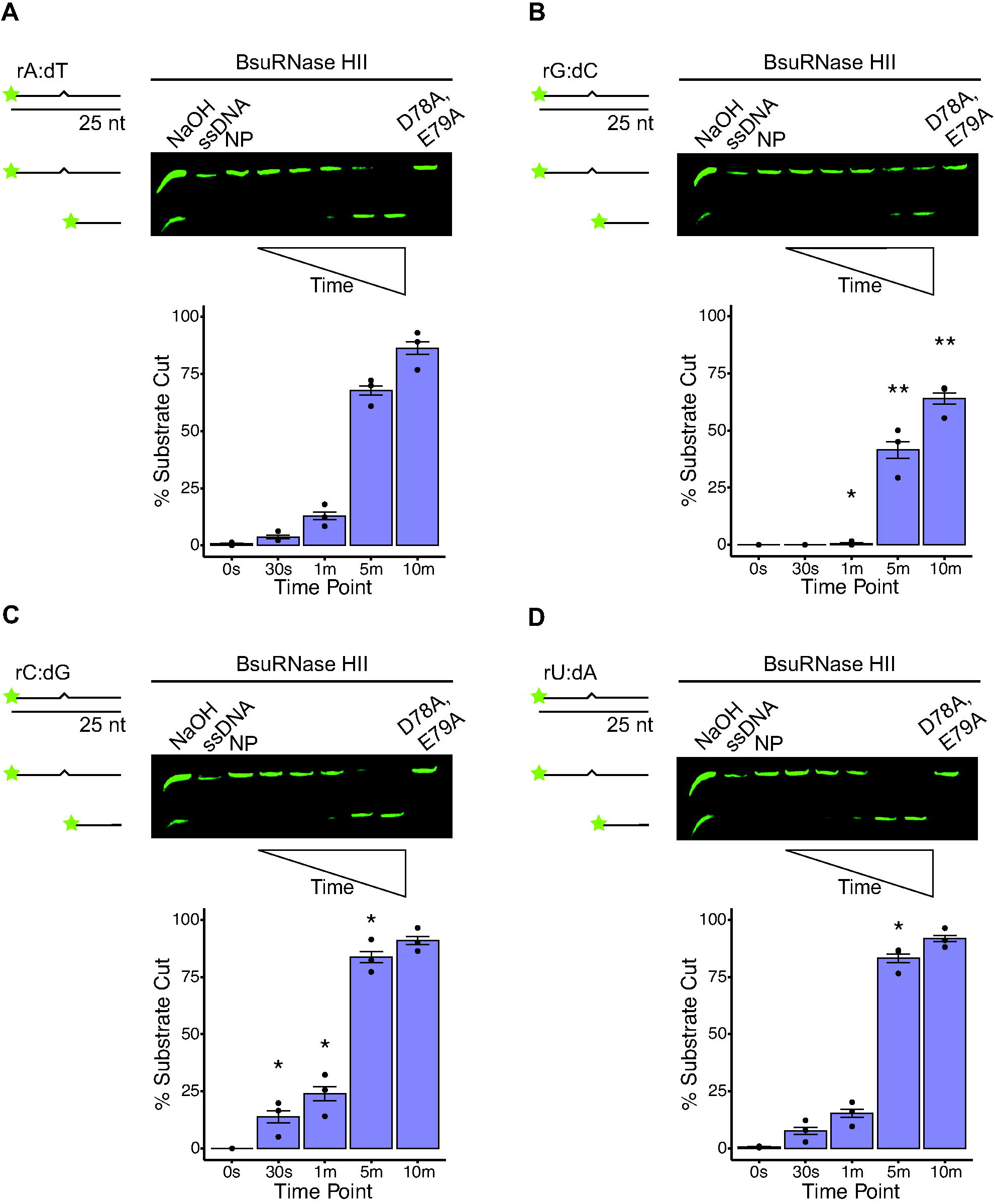
BsuRNase HII demonstrates weaker nuclease activity on rG:dC relative to other rNMPs. Representative urea-PAGE for assays performed over 10 minutes at 37°C with 50 nM *B. subtilis* RNase HII and 100 nM of the following 25-nucleotide substrates: (A) rA:dT, (B) rG:dC, (C) rC:dG, and (D) rU:dA. Each substrate is composed of 25-nucleotide long dsDNA with a single rNMP, represented by a zigzag. Quantification below each gel shows the mean percent of substrate incised over time (0 s, 30 s, 1 min., 5 min., and 10 min. time points) with black bars to show the standard error, for three replicates. Asterisks indicate significance, with * representing p < 0.05, ** representing p < 0.01, and *** representing p < 0.001 as compared to the same time point for rA:dT. Each gel contains ssDNA, no protein (NP), and catalytic impaired (D78A, E79A) controls. An alkaline ladder, represented by “NaOH”, was prepared by incubating substrate with 200 nM final NaOH. rA:dT was prepared with oJC3 and oJC2, rG:dC was prepared with oJC3 and oJC5, rC:dG was prepared with oJC6 and oJC7, and rU:dA was prepared with oJC8 and oJC9.

### BsuRNase HII inefficiently incises mismatched rGMP

Because rNMPs are the most frequent replication error made *in vivo*, it stands to reason that rNMPs could be mismatched, resulting in an error that could be recognized by RNase HII or MutS during MMR (Shen *et al*., 2011). Since dA:dC and dG:dT are the two most common mismatches formed *in vivo* (Kunkel, 2004, Kunkel & Erie, 2005, Lahue *et al*., 1989, Modrich & Lahue, 1996), we tested BsuRNase HII on the following rNMP-containing mismatched substrates: rA:dC and rG:dT. We found that BsuRNase HII incised rA:dC as efficiently as the properly base-paired ribonucleotides, suggesting that rA:dC could serve as a substrate *in vivo* **(Figure 3)**. Consistent with our previous observation for rGMP in a Watson-Crick base pair, BsuRNase HII exhibits poor activity on mismatched rGMP (rG:dT) We compared the 10 min time point of the matched (rG:dC) to mismatched rG:dt and found a significant decrease in processing. Thus even though matched rGMP is processed inefficiently, mismatched rGMP is significantly worse **(Figure 3B)**. We used both a no protein control and catalytically-impaired variant as a negative control, in addition to generating an alkaline ladder with substrate as described earlier. Moreover, we also incubated BsuRNase HII in lane 2 with a canonical rA:dT base pair as an additional control to show the protein was active **(Figure 3)**. Contrary to our observation for BsuRNase HII, we show that EcoRNase HII efficiently incises both mismatched ribonucleotides to nearly the same extent as properly base-paired control ribonucleotides any perceived difference between the two mismatched substrates was not statistically significant **(Figure 3C and D)**. We conclude that BsuRNase HII recognizes mismatched rAMP but is poor at recognition of mismatched rGMP **(see Discussion)**.

**Figure 3.**
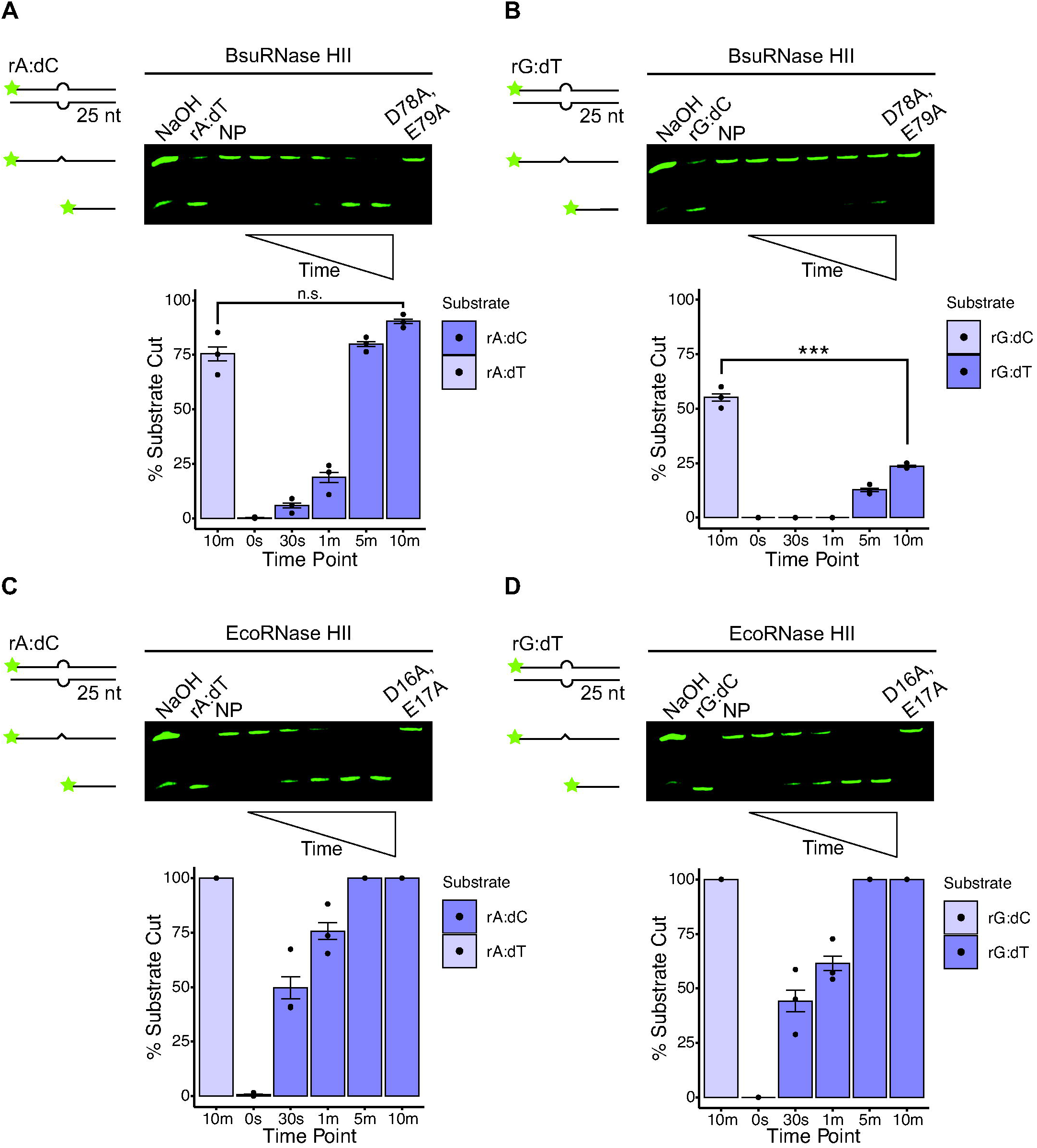
EcoRNase HII is more active on mismatched rNMPs than BsuRNase HII. Representative urea-PAGE for assays performed over 10 minutes at 37 °C with (A) 50 nM BsuRNase HII and 100 nM rA:dC, (B) 50 nM BsuRNase HII and 100 nM rG:dT, (C) 6.25 nM EcoRNase HII and 100 nM rA:dC, and (D) 6.25 nM EcoRNase HII and 100 nM rG:dT. Each substrate is composed of 25-nucleotide long dsDNA with a single rNMP forming a mismatched base pair, represented by a bulge. Quantification below each gel shows the mean percent of substrate incised over time (0 s, 30 s, 1 min., 5 min., and 10 min. time points) with black bars to show the standard error, for three replicates. Asterisks indicate significance, with * representing p < 0.05, ** representing p < 0.01, and *** representing p < 0.001. Each gel contains positive (canonical base pair), no protein (NP), and catalytic impaired controls. An alkaline ladder, represented by “NaOH”, was prepared by incubating substrate with 200 nM final NaOH. rA:dC was prepared with oJC3 and oJC5, and rG:dT was prepared with oJC4 and oJC2.

### BsuRNase HII is refractory to damaged ribonucleotides *in vitro*

It has been shown previously that EcoRNase HII can incise damaged ribonucleotides, while eukaryotic RNase H2 orthologs cannot (Malfatti *et al*., 2019). We tested BsuRNase HII on r8oG:dC and an abasic site with a ribose sugar, denoted as rOH:dC. We show that BsuRNase HII is refractory to incise both damaged ribonucleotides **(Figure 4A and B)**. Additionally, upon longer incubation of BsuRNase HII with r8oG:dC, we still do not observe incision **(Figure 4E)**. As a control, we show that BsuRNase HII cleaves an undamaged rNMP-containing substrate (rG:dC) **(Figure 4A and B)**. Alkaline ladder was generated with damaged substrate as described above. We also tested EcoRNase HII on the same damaged substrates. We did not observe activity on the ribose-containing abasic site (rOH:dC), although we observed some activity on the r8oG:dC **(Figure 4C and D)**. Our results differ slightly from a prior report, which observed weak EcoRNase HII activity on rOH (Malfatti *et al*., 2017). Although our results are not identical, we do observe a similar trend in that r8oG serves as a reasonable substrate for EcoRNase HII, while rOH does not. We further tested BsuRNase HII and EcoRNase HII on r8oG using a 2-fold increase in Mg^2+^ to 2 mM, as well as 1 mM Mn^2+^. We did not observe BsuRNase HII incision of r8oG under either metal condition, while EcoRNase HII cleaved r8oxoG under both metal conditions **(Figure S1).** With our work, we conclude that BsuRNase HII is unable to incise damaged ribonucleotides, while EcoRNase HII can repair r8oG and not rOH (abasic). Further, with these results, we suggest that rOH sites would need to be repaired through BER (Wozniak & Simmons, 2022). We conclude that rOH is either not repaired or inefficiently repaired by RER in bacteria.

**Figure 4.**
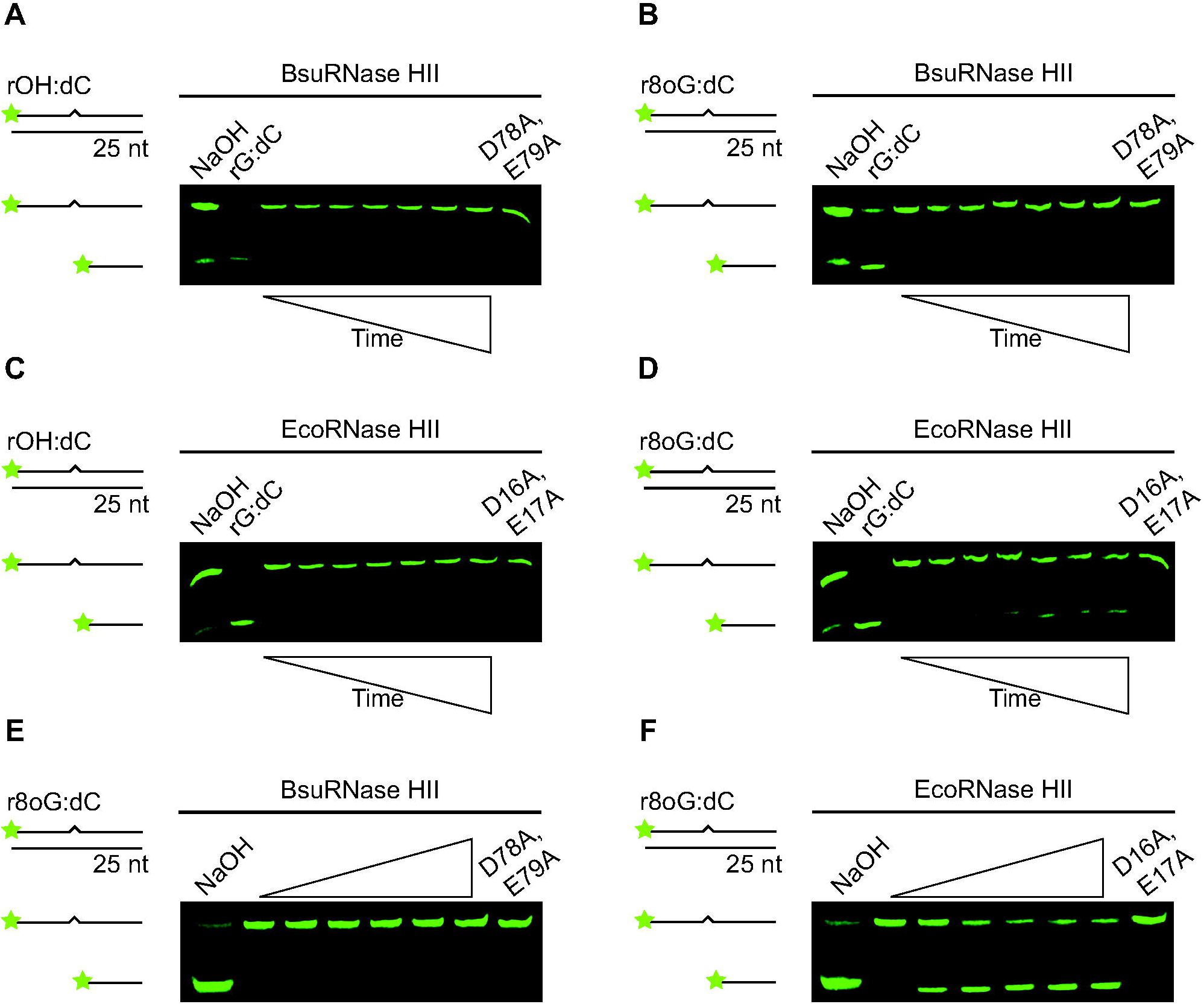
EcoRNase HII is more active on damaged rNMPs than BsuRNase HII. Representative urea-PAGE for assays performed in triplicate over 160 minutes at 37°C. r8oxoG is abbreviated r8oG. The following conditions were used in (A) 50 nM BsuRNase HII and 100 nM rOH:dC, (B) 50 nM BsuRNase HII and 100 nM r8oG:dC, (C) 6.25 nM EcoRNase HII and 100 nM rOH:dC, and (D) 6.25 nM EcoRNase HII and 100 nM r8oG:dC. Each substrate is composed of 20-nucleotide long dsDNA with a single rNMP, represented by a zigzag. In the case of rOH:dC, the sugar is intact, but the base is missing (abasic). The time points for all four gels are the same: 0 s, 5 min., 10 min., 20 min., 40 min., 80 min., and 160 min. Each gel contains positive (rG:dC) and catalytic impaired controls. An alkaline ladder, represented by “NaOH”, was prepared by incubating substrate with 200 nM final NaOH. rOH:dC was prepared with oJC10 and oJC5, and r8oG:dC was prepared with oJC11 and oJC5.

## DISCUSSION

It is well established that single ribonucleotide errors represent the most frequent nucleotide in need of repair (Williams *et al*., 2016, Zhou *et al*., 2021, Schroeder *et al*., 2015). Given that replicative and non-replicative DNA polymerases make sugar errors at such a high frequency (Nick McElhinny *et al*., 2010b, Yao *et al*., 2013), it would be expected that mismatched and damaged rNMPs would be present in the genome and require repair. The increased likelihood of mismatched and damaged rNMPs in genomic DNA raises the question of how these complex errors are repaired. Our results show that bacterial RNase HII efficiently resolves mismatched rNMPs, while the activity observed for canonical and damaged rNMPs depends on the specific bacterial enzyme tested. These observations suggest that RNase HII substrate preference varies within the bacterial kingdom. This divergence in activity is especially interesting, as RNase HII is the only conserved RNase H family member between *E. coli* and *B. subtilis* (Ohtani *et al*., 1999b).

The crystal structure of human RNase H2 shows that both an invariant tyrosine residue and a glycine-arginine-glycine (GRG) motif, together called the junction-sensing module, form hydrogen bonds with the 2′-OH of the ribose sugar (Chon *et al*., 2006, Lai *et al*., 2000). This module sets RNase HII apart from other family enzymes like RNase HI or RNase HIII, which lack the central basic residue of the GRG motif and is thus unable to precisely recognize a single RNA-DNA junction (Nowotny *et al*., 2005, Nowotny & Yang, 2006, Hyjek *et al*., 2019). Given the structural data, it would seem that RNase HII would only require a 2′-OH and an RNA-DNA junction to recognize all possible ribonucleotide variants (Chon *et al*., 2006, Lai *et al*., 2000). In most prior studies, a single ribonucleotide has been chosen and studied as representative of all ribonucleotides cleaved by RNase HII. Given that each of the four rNMPs are incorporated into DNA at different rates (Yao *et al*., 2013, Balachander *et al*., 2020), we asked if bacterial RNase HII incises the four canonical rNMPs with similar efficiencies. We found that EcoRNase HII does indeed incise all four canonical rNMPs with similar efficiencies. Incision of rCMP is the least efficient, but overall rAMP, rGMP, rCMP and rUMP evoke similar processing by EcoRNase HII. Based on *in vitro* rNMP incorporation numbers, we would expect roughly 1533 rAMP, 331 rCMP, 124 rGMP and 6 rUMP per round of replication for *E. coli* DNA polymerase III holoenzyme (Yao *et al*., 2013). Thus, even though rAMP is by far the most frequently incorporated rNMP by the *E. coli* replicase, EcoRNase HII incises all rNMPs equally well. In the case of BsuRNase HII, we consistently observe inefficient processing of rGMP, with rAMP, rCMP, and rUMP being processed with similar efficiencies. The reason for slow incision of rGMP by BsuRNase HII is unclear. *B. subtilis* is a low-GC organism, although BsuRNase HII efficiently cleaved rCMP and rUMP, suggesting that the difference is not shaped by the abundance of particular rNMPs in the genome. Given that BsuRNase HII is so slow to process rGMP, we speculate that rGMP may persist *in vivo,* potentially leading to mutations and genome instability.

An important finding from our work is that both *E. coli* and *B. subtilis* RNase HII process rUMP efficiently, which is impressive given that we expect about 5-6 rUMP misincorporations per genome replication event (Yao *et al*., 2013). Although the number of rUMPs incorporated into the genome is expected to be low, another source of rUMP formation could be through deamination of rCMP to rUMP. Therefore, one possibility is that rUMP is higher in abundance in the genome than anticipated because of both sugar errors by replicative DNA polymerases and deamination events. Bacterial UDG is adept at removing dUMP from dsDNA and ssDNA, but it cannot use an ssRNA substrate (Pearl, 2000, Savva *et al*., 1995). A recent paper found that UDG can remove a single rUMP from DNA *in vitro* however, it does not appear to do so *in vivo* (Fan *et al*., 2025). Given that UDG does not efficiently use an RNA substrate due to clash of a conserved phenylalanine residue with the 2′-OH (Savva *et al*., 1995), rUMP would need to be repaired exclusively through RER. This would place RNase HII in an important role for removing rUMP formed through both DNA polymerase misincorporation events and deamination of rCMP to rUMP. Further, since uracil is mutagenic and results in GC-to-AT transitions (Duncan & Miller, 1980, Duncan *et al*., 1978), RNase HII would help ensure fidelity during the replication process by removing rUMP.

In the case of mismatched rNMPs, we tested rA:dC and rG:dT because these ribonucleotide mismatches should be the most commonly mismatched rNMPs, if rNMP errors follow the same trend as dNMP errors during replication (Kunkel, 2004, Kunkel & Bebenek, 2000, Kunkel & Erie, 2005). EcoRNase HII is able to recognize both mismatches equally well, while BsuRNase HII efficiently incises rA:dC but not rG:dT. The fact that BsuRNase HII struggles to process rG:dT in a mismatch provides further validation that the *B. subtilis* enzyme is inefficient at processing rGMP. Prior work showed that EcoRNase HII was more active on mismatched rather than matched rNMPs (Malfatti *et al*., 2019). We also find that EcoRNase HII processes mismatched rNMPs more rapidly than those that are properly base-paired. We speculate that when rAMP is involved in a mismatch, the rA:dC mispair may provide RNase HII with better recognition of the 2′-OH. Given that bacterial RNase HII recognizes mismatched ribonucleotides quite well, we find it interesting that eukaryotic RNase H2 does not (Malfatti *et al*., 2019). Therefore, we suggest that in eukaryotes, MMR addresses mismatched rNMPs, while bacterial MMR machinery is either poor at recognizing mismatched rNMPs or RER and MMR cooperate to correct mismatched rNMPs in the genome. Since bacterial MutS has not been tested on rNMP-containing mismatches, we speculate that mismatched rNMPs are repaired by MMR and RER, providing two avenues for their removal **(Figure 5)**.

**Figure 5.**
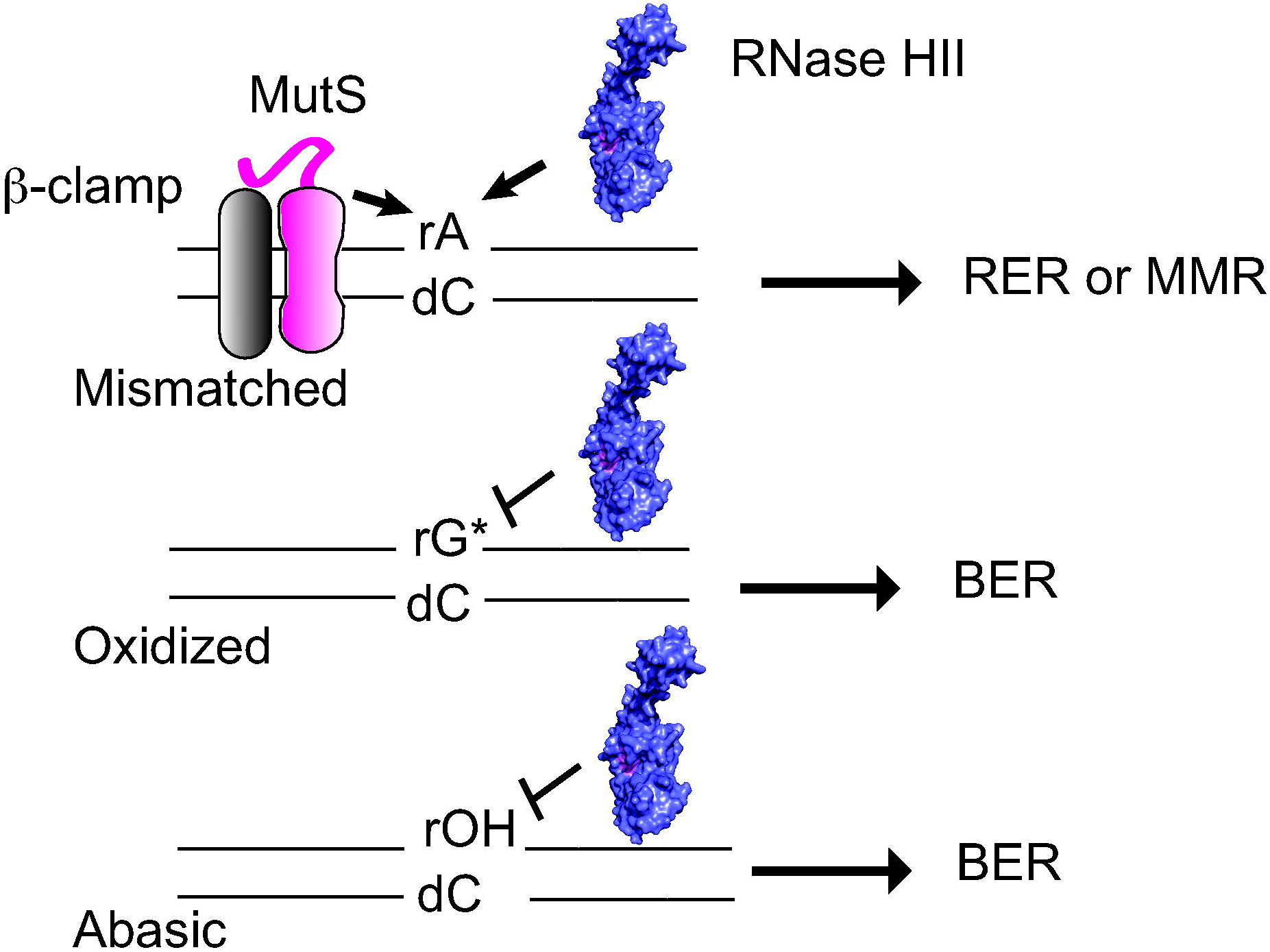
BsuRNase HII and mismatch repair correct rNMP mismatches while BER repairs damaged rNMPs. Shown is a model for the repair of mismatched and damaged rNMPs in *B. subtilis*. MutS with the aid of β-clamp (DnaN) and RNase HII can bind and correct rNMP errors. Since RNase HII is inefficient with rG:dT mismatches we propose that MutS corrects rG:dT while both MutS and RNase HII correct rA:dC and other mispairs. Since BsuRNase HII is refractory to r8oG (rG*) and rOH, we propose that both are repaired through BER pathway.

In the case of damaged rNMPs, we provide clear evidence that BsuRNase HII does not repair rOH or r8oG. We also show that EcoRNase HII repairs r8oG but not rOH. Eukaryotic RNase H2 and archaeal RNase H2 are refractory to incision of rOH and r8oG (Malfatti *et al*., 2017, Malfatti *et al*., 2019), and it has been suggested that recognition of damaged rNMPs has been lost through evolution from bacteria to eukaryotes (Malfatti *et al*., 2019). Like dNMPs, rNMPs are subject to oxidative damage and other modifications (Randerath *et al*., 1992, Loeb & Preston, 1986). Our results show that the capacity of bacterial RNase HIIs to repair damaged rNMPs varies between *B. subtilis* and *E. coli* two organisms separated by approximately two billion years of evolution (Pace *et al*., 1986). The ability of RNase H enzymes to recognize and repair damaged ribonucleotides may simply reflect the DNA repair repertoire of each organism tested. In the case of eukaryotes, AP endonuclease 1 (APE1) removes rOH and r8oG (Malfatti *et al*., 2017). Since APE1 is so efficient, there is no need for RNase H2 to recognize these lesions. For *E. coli* and *B. subtilis*, the role of RNase HII in repair of rOH and oxidative damage is less clear. EcoRNase HII can incise r8oG well, suggesting that EcoRNase HII and MutY may overlap to repair r8oG. Under the conditions tested here, EcoRNase HII and BsuRNase HII do not process rOH lesions, while BsuRNase HII does not repair r8oG. Based on our evidence we conclude that rOH sites in both organisms and r8oG in *B. subtilis* would be repaired by BER **(Figure 5)**.

## EXPERIMENTAL PROCEDURES

### Plasmid Construction

A list of strains, plasmids, and primers used in this work is provided in **Table S1**, **Table S2,** and **Table S3** respectively. The *B. subtilis* PY79 *rnhB* gene (oJR88 and oJR89) and pE-SUMO backbone (oJR46 and oJR47) were PCR amplified from purified JRC9 gDNA and Hind III-digested pFCL1, respectively. These fragments were joined using Gibson Assembly to make pFCL25. Site-directed mutagenesis of pFCL25 (oJR94 and oJR95) was performed to generate pFCL26. Similarly, the *E. coli* MC1061 *rnhB* gene (prJC36 and prJC40) and pE-SUMO backbone (prJC44 and prJC45) were PCR amplified from purified JRC6 gDNA and HindIII-digested pJC1, respectively. The fragments were joined using the Gibson Assembly to generate pJC10, which was subjected to site-directed mutagenesis (prJC41 and prJC42) to make pJC11. The assembled plasmids were used to transform chemically competent *E. coli* MC1061, which was plated on LB agar with 25 µg/mL kanamycin (kan). Kanamycin-resistant colonies were subject to colony PCR (*B. subtilis*: oJR88 and oJR89, *E. coli*: prJC36 and prFCL25), and the plasmid sequences were verified via Sanger sequencing (*B. subtilis*: prFCL25, *E. coli*: T7 and T7term) using Eurofins Scientific.

### Protein Expression

Each expression plasmid was purified from the appropriate *E. coli* MC1061 strain and subsequently used to transform competent *E. coli* BL21(DE3). Transformants were plated on LB agar (25 µg/mL kan) and incubated at 37°C overnight. A single colony was picked to inoculate 50 mL LB (25 µg/mL kan) and incubated at 37°C overnight while shaking at 200 rpm. 5-15 mL of this overnight culture were used to inoculate 1 L of LB (25 µg/mL kan) under sterile conditions. Scaled-up culture was grown at 37°C and 165-200 rpm until the OD_600_ reached between 0.5 and 0.7. Expression was induced with 0.5 mM isopropyl-β-D-thiogalactopyranoside (IPTG) for 3 hours at 37°C between 165-200 pm. Cells were harvested by centrifugation at 4000 rpm and 4°C for 30 minutes, after which each pellet was resuspended in 50 mL LB. Resuspended pellets were spun down again at 4000 rpm and 4°C for 30 minutes. Once the supernatant was decanted, the pellets were weighed, frozen, and then stored at –80°C until purification.

### Protein Purification

Pellets were thawed on ice and resuspended in 30 mL Nickel Buffer A (20 mM Tris-HCl pH 8.0, 400 mM NaCl, 5% glycerol, and 1 mM DTT) using a serological pipet. After the edition of 1 EDTA-free Pierce Protease Inhibitor tablet per pellet, the combined pellets were sonicated at 75% amplitude for an “on time” of 1 minute and 40 seconds and spun down at 30,000 x g and 4°C for 40 minutes. Clarified lysate was loaded onto an equilibrated Ni-NTA affinity column with a 5-mL column volume (CV) and the flow-through (FT) was collected. The column was washed with 5 CV of 100% Nickel Buffer A and 10 CV of 5% Nickel Buffer B (20 mM Tris-HCl pH 8.0, 400 mM NaCl, 5% glycerol, 1 mM DTT, and 300 mM imidazole); both washes were collected. The His_6_-tagged protein was eluted in 1-mL fractions using 16 mL of 100% Nickel Buffer B. The concentration of each fraction was measured on a NanoDrop Microvolume Spectrophotometer, and those with an A_280_ close to or greater than 1 were pooled. Pooled fractions were diluted using Dilution Buffer (20 mM Tris-HCl pH 8.0, 5% glycerol, 1 mM EDTA, and 1 mM DTT) to reduce the salt to 300 mM and supplemented with 1 mM DTT. The pooled protein was then incubated with purified SUMO protease for 2 hours at room temperature. Post-incubation, the protein was carefully pipetted into 10,000 MWCO dialysis tubing and dialyzed overnight at 4°C against Dialysis Buffer (20 mM Tris-HCl pH 8.0, 300 mM NaCl, and 5% glycerol) to remove imidazole and DTT.

The following morning, the dialyzed protein was filter-sterilized using a 50-mL syringe and a 0.22-µM sterile filter that had been primed with Dialysis Buffer. The filter-sterilized protein was added to the Ni-NTA affinity column, which had been equilibrated with Dialysis Buffer and stored the day before, and the FT was collected. Subsequently, 3 CV of filtered Dialysis Buffer were added to the column and collected as the Dialysis Wash. Finally, 3 CV of 5% Nickel Buffer B and 100% Nickel Buffer B were added sequentially to remove residual His_6_-tag and SUMO protease from the resin. SDS-PAGE using a 10% polyacrylamide gel was used to verify the presence of pure protein in the filter-sterilized protein FT and Dialysis Wash. These fractions were combined and concentrated using a 10,000 MWCO Amicon Ultra Centrifugal Filter. Glycerol was added to concentrated protein to 25% and the protein was flash frozen using liquid nitrogen before storing at –80°C.

Additional purification was performed using an ÄKTA FPLC. Both crude BsuRNase HII and EcoRNase HII were subjected to anion exchange using a HiTrap^TM^ Q FF column (Cytiva 17505301) in Buffer A (20 mM Tris pH 8.0, 1 mM DTT, and 5% glycerol) and Buffer B (20 mM Tris pH 8.0, 1 mM DTT, 5% glycerol, and 500 mM NaCl). Sample, diluted to 50 mM NaCl in IEC Buffer A, was injected onto the column using a 50-mL Superloop^TM^ (Cytiva 18111382). The column was equilibrated with 10% IEC Buffer B, and protein was eluted via fractionation over a gradient of 100% IEC Buffer B. Peak fractions were identified using SDS-PAGE, pooled, and concentrated using a 10,000 MWCO Amicon Ultra Centrifugal filter. EcoRNase HII and EcoRNase HII D16A, E17A were combined with glycerol to 25% and flash frozen using liquid nitrogen before storing at –80°C.

BsuRNase HII and BsuRNase HII D78A, E79A were also purified via size exclusion chromatography using a HiPrep Sephacryl^TM^ S-200 HR preparative column (Cytiva 17116601) and Buffer A (20 mM Tris pH 8.0, final NaCl concentration of pooled protein from anion exchange, and 1 mM DTT). The column was equilibrated with Buffer A, and protein was eluted via fractionation. SDS-PAGE was used to determine the peak fractions, which were then combined and concentrated as described above. Glycerol was added to 25%, and protein was flash frozen using liquid nitrogen before storing at – 80°C. A representative 10% SDS-PAGE gel with 1 ug of each purified protein and stained with Coomassie blue dye is included **(Figure S2).**

### RNase HII activity assays

Assays performed herein were based upon a previously established method (Randall *et al*., 2019, Schroeder *et al*., 2023, Schroeder *et al*., 2017, Lowder & Simmons, 2023). RNA-DNA hybrid substrates were prepared by combining 1 μM 5′-labeled rNMP-containing oligonucleotide with 2 μM unlabeled complement in 1X TS Buffer (20 mM Tris pH 8.0, 100 mM NaCl), boiling at 98°C for 1 min., and cooling to anneal away from light. Oligonucleotides were used to assemble the following substrates: single canonical rNMP-containing hybrids (oJC3, oJC4, oJC6, and oJC8), and their complements (oJC2, oJC5, oJC7, and oJC9), single mismatched rNMP-containing hybrids (oJC3:oJC5 and oJC4:oJC2), and single damaged rNMP-containing hybrids (oJC10:oJC5 and oJC11:oJC5). All oligonucleotides were 25 nucleotides long, and their sequences can be found in **Table S4**. Purified protein was diluted to a working concentration of 200 or 100 nM in 1X MgTS Buffer (20 mM Tris pH 8.0, 100 mM NaCl, 1 mM MgCl_2_). Reactions were prepared in 1X MgTS Buffer using 50 nM BsuRNase HII or 6.25 nM EcoRNase HII and 100 nM annealed substrate. Reactions were incubated on a heat block at 37°C, during which 8 µL time points were taken (0 s, 30s, 1 min., 5 min., 10 min., unless otherwise noted), mixed 1:1 with 2X stop buffer (95% formamide, 20 mM EDTA, 0.01% bromophenol blue), boiled at 98°C for 5 min., and immediately stored on ice. The 0s time point was prepared before starting the reaction by mixing diluted protein and reaction buffer in the appropriate ratio and immediately mixing with 2X stop buffer. Appropriate controls were performed depending on the nature of the assay, including no protein, catalytic impaired, ssDNA containing an rNMP, and/or a canonical hybrid substrate (in the case of the damaged rNMP assays, rG:dC). The ladder was prepared by incubating substrate with an equivalent amount of 400 nM NaOH at 37°C for 10 min., followed by boiling. Products were electrophoresed using 20% denaturing urea-PAGE (8M urea) and imaged using the 800 nm channel of a LiCor Odyssey imager. Three replicates were performed for each substrate.

### Activity assay analysis

Analysis for activity assays has been previously described (Lowder & Simmons, 2023, Randall *et al*., 2017, Schroeder *et al*., 2017). FIJI was used for the quantification of gel band intensities. The time point lanes were selected, using the 0s lane as the reference. The intensities corresponding to each lane were plotted, and the Wand Tool was used to determine the area under the curve for each lane. Measurements from three replicates were exported to Microsoft Excel and used to determine the relative percent of cut substrate (area under curve for time point divided by area under curve for 0-second time point, times 100). The following packages were used to process the data and generate graphs in R: *tidyverse*, (*ggplot2*), and *forcats* (Wickham, 2016, Wickham H *et al*., 2019).

### Statistical analysis

Statistical analyses were performed in R using the base package (RC, 2022). Using the lm() function, the data were fit to an ordinary least squares (OLS) linear regression model, where percent incision was treated as the response and substrate as the sole predictor. A separate model was generated for substrates at each time point (i.e., 10 minutes, etc.). For the BsuRNase HII assays with canonical rNMPs, all comparisons were made with rA:dT as the reference. Similarly, for BsuRNase HII assays with rA:dC and rG:dT, comparisons were made at 10 minutes only with rA:dT and rG:dC as the references, respectively. Using an alpha (17) of 0.05, significance is shown in each figure with * representing p < 0.05, ** representing p < 0.01, and *** representing p < 0.001.

## ACKNOWLEDGEMENTS

This work was funded by National Institutes of Health grant R35GM131772 to LAS. We thank Caroline Lowder for building the *rnhB* expression vector. JRC was supported by the following NIH training grant: Michigan Predoctoral Training in Genetics (T32GM007544). JRC was also supported by precandidate and candidate research grants from the Horace H. Rackham Graduate School, in addition to two Fellowships (Peter Olaus Okkelberg and Edwin H. Edwards) from the Department of Molecular, Cellular, and Developmental Biology at the University of Michigan. We thank the Nandakumar, Bardwell, and Jakob labs for use of equipment during this project.

## DATA AVAILABILITY

All data relevant to this manuscript are reported in the main or supporting information. Any other information is available upon request.

## AUTHOR CONTRIBUTIONS

Julianna R Cresti Conceptualization, Formal analysis, Funding acquisition, Investigation, Validation, Visualization, Writing original draft, Writing-review and editing.

Lyle A. Simmons Conceptualization, Formal analysis, Funding acquisition, Investigation, Project administration, Supervision, Validation, Visualization, Writing original draft, Writing-review and editing.

## SUPPLEMENTAL MATERIAL

**Supporting Table S1**. Strains used in this study

**Supporting Table S2.** Plasmids used in this study.

**Supporting Table S3**. Oligonucleotide primers used in this study.

**Supporting Table S4**. Oligonucleotides used in this study

**Figure S1. Changing metal and Mg^2+^ concentration does not affect BsuRNase HII nuclease activity on r8oG:dC.** Shown are representative urea-PAGE assays performed over a 160-minute time course at 37°C with (A) 50 nM *B. subtilis* RNase HII and 100 nM r8oG:dC, (B) 50 nM *B. subtilis* RNase HII and 100 nM r8oG:dC, (C) 6.25 nM *E. coli* RNase HII and 100 nM r8oG:dC, and (D) 6.25 nM *E. coli* RNase HII and 100 nM r8oG:dC. Assays shown in (A) and (C) were performed with 2 mM MgCl_2_, whereas(B) and (D) were performed with 1 mM MnCl_2_. Each substrate is composed of a 25-oligonucleotide dsDNA with a single rNMP, represented by a zigzag. Each gel contains a canonical base pair (rG:dC) and catalytic impaired RNase HII controls. An alkaline ladder was prepared by incubating each substrate with 200 nM NaOH. rOH:dC was prepared with oJC10 and oJC5, and r8oG:dC was prepared with oJC11 and oJC5.

**Figure S2. Purified BsuRNase HII and EcoRNase HII.** Shown is an SDS-PAGE. Lane 1 contains a protein dual color standards with the corresponding molecular weight (MW) indicated. Lanes 2-5 each contain 1 µg of *B. subtilis* RNase HII (28.4 kDa), *B. subtilis* RNase HII D78A E79A (28.3 kDa), *E. coli* RNase HII (21.5 kDa), and *E. coli* RNase HII D16A E17A (21.4 kDa), respectively.

