## Supporting tables and figures for "*Bacillus subtilis* RNase HII is inefficient at processing guanosine monophosphate and damaged ribonucleotides"

**Short title:** RNase HII resolves canonical and mismatched ribonucleotides

**Keywords:** RNase HII, *Bacillus subtilis*, DNA replication, DNA damage, ribonucleotides, mismatches

**Supporting Table S1**. Strains used in this study.

| Strain ID | Species | Strain | Genotype/Plasmid | Reference |
|---|---|---|---|---|
| FCL1 | *E. coli* | MC1061 | pFLC1 | |
| FCL70 | *E. coli* | MC1061 | pFCL25 | |
| FCL71 | *E. coli* | BL21(DE3) | pFCL25 | |
| FCL72 | *E. coli* | MC1061 | pFCL26 | |
| FCL73 | *E. coli* | BL21(DE3) | pFCL26 | |
| JRC5 | *E. coli* | BL21(DE3) | | |
| JRC6 | *E. coli* | MC1061 | | |
| JRC9 | *B. subtilis* | PY79 | Prototroph Spβ° | |
| JRC20 | *E. coli* | MC1061 | pJC10 | |
| JRC21 | *E. coli* | BL21(DE3) | pJC10 | |
| JRC22 | *E. coli* | MC1061 | pJC11 | |
| JRC23 | *E. coli* | BL21(DE3) | pJC11 | |

**Supporting Table S2.** Plasmids used in this study.

| Name | Genotype | Reference |
|---|---|---|
| pFCL1 | $His_6$-SUMO-*B. subtilis fenA* | |
| pFCL25 | $His_6$-SUMO-*B. subtilis rnhB* | |
| pFCL26 | $His_6$-SUMO-*B. subtilis rnhB* D78A E79A | |
| pJC10 | $His_6$-SUMO-*E. coli rnhB* | |
| pJC11 | $His_6$-SUMO-*E. coli rnhB* D16A E17A | |

**Supporting Table S3**. Oligonucleotides used as primers in this study.

| Oligo | Sequence |
|---|---|
| **oJR46** | tcgagcaccaccaccaccaccactgag |
| **oJR47** | acctccaatctgttcgcggtgagcctcaataatatcg |
| **oJR88** | ccgcgaacagattggaggtgtgaatacattaaccgtaaaggacattaaagacc |
| **oJR89** | tggtggtggtggtgctcgattatctgaaagattgaacaggagcg |
| **oJR92** | gtgaaatacttgggctgacagactc |
| **oJR94** | gttgccgcggtcggccg |
| **oJR95** | gaccgcggcaacacctgcaatc |
| **prFCL25** | cctttcgggctttgttagcagcc |
| **prJC36** | accgcgaacagattggaggtatgatcgaatttgtttatcc |
| **prJC40** | gtggtggtggtggtgctcgatcaggacgcaagtc |
| **prJC41** | tgtggctgcagtcggacgcgggccgttagttg |
| **prJC42** | ccgactgcagccacacccgcaaccagctgcg |
| **prJC44** | acctccaatctgttcgcggtg |
| **prJC45** | tcgagcaccaccaccaccac |
| **prJC51** | tccggcgtagaggatcgagatctcg |
| **prJC52** | cgggctttgttagcagccggatctc |

Primers were used for PCR amplification, mutagenesis, Gibson assembly, and colony PCR.

**Supporting Table S4**. Oligonucleotides used as substrates for RNase HII.

| Oligo | Sequence | Label |
|---|---|---|
| **oJC1** | ggcttatacagcatcgagctcagga | 5' IR800CWN |
| **oJC2** | tcctgagctcgatgctgtataagcc | n/a |
| **oJC3** | ggcttatacagcrAtcgagctcagga | 5' IR800CWN |
| **oJC4** | ggcttatacagcrGtcgagctcagga | 5' IR800CWN |
| **oJC5** | tcctgagctcgacgctgtataagcc | n/a |
| **oJC6** | ggcttatacagcrCtcgagctcagga | 5' IR800CWN |
| **oJC7** | tcctgagctcgaggctgtataagcc | n/a |
| **oJC8** | ggcttatacagcrUtcgagctcagga | 5' IR800CWN |
| **oJC9** | tcctgagctcgaagctgtataagcc | n/a |
| **oJC10** | ggcttatacagcrOHtcgagctcagga | 5' IR800CWN |
| **oJC11** | ggcttatacagcr8-oGtcgagctcagga | 5' IR800CWN |

Oligos were used to assemble double-stranded substrates for RNase HII endonuclease assays. All oligos are 25 nucleotides in length. The single rNMP modification has been underlined for clarity.

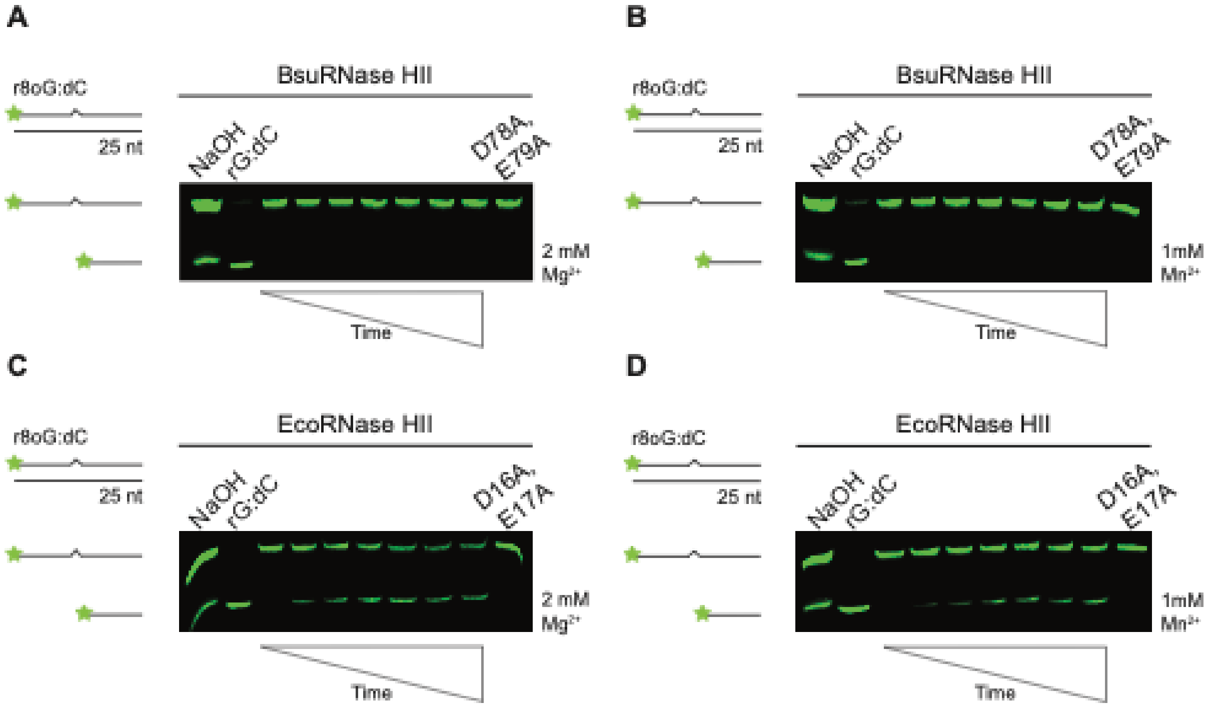


**Figure S1**. **Changing metal and $Mg^{2+}$ concentration does not affect BsuRNase HII nuclease activity on r8oG:dC.** Shown are representative urea-PAGE assays performed over a 160-minute time course at 37ºC with (A) 50 nM *B. subtilis* RNase HII and 100 nM r8oG:dC, (B) 50 nM *B. subtilis* RNase HII and 100 nM r8oG:dC, (C) 6.25 nM *E. coli* RNase HII and 100 nM r8oG:dC, and (D) 6.25 nM *E. coli* RNase HII and 100 nM r8oG:dC. Assays shown in (A) and (C) were performed with 2 mM $MgCl_2$, whereas (B) and (D) were performed with 1 mM $MnCl_2$. Each substrate is composed of a 25-oligonucleotide dsDNA with a single rNMP, represented by a zigzag. Each gel contains a canonical base pair (rG:dC) and catalytic impaired RNase HII controls. An alkaline ladder was prepared by incubating each substrate with 200 nM NaOH. rOH:dC was prepared with oJC10 and oJC5, and r8oG:dC was prepared with oJC11 and oJC5.

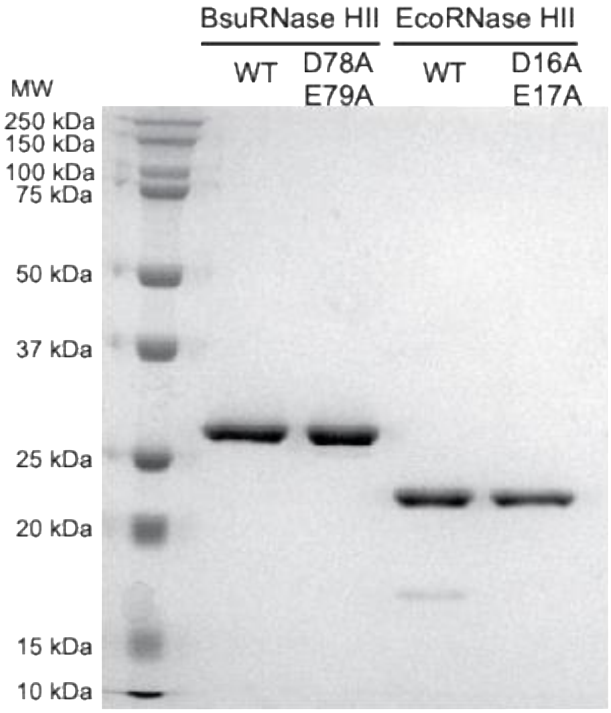


**Figure S2. Purified BsuRNase HII and EcoRNase HII.** Shown is an SDS-PAGE. Lane 1 contains a protein dual color standards with the corresponding molecular weight (MW) indicated. Lanes 2-5 each contain 1 µg of *B. subtilis* RNase HII (28.4 kDa), *B. subtilis* RNase HII D78A E79A (28.3 kDa), *E. coli* RNase HII (21.5 kDa), and *E. coli* RNase HII D16A E17A (21.4 kDa), respectively.

### REFERENCES
